# Effects of perfluoroalkyl and polyfluoroalkyl substances on soil structure and function

**DOI:** 10.1101/2021.10.26.465889

**Authors:** Baile Xu, Gaowen Yang, Anika Lehmann, Sebastian Riedel, Matthias C. Rillig

## Abstract

Soils are impacted at a global scale by several anthropogenic factors, including chemical pollutants. Among those, perfluoroalkyl and polyfluoroalkyl substances (PFAS) are of concern due to their high environmental persistence, and as they might affect soil health and functions. However, data on impacts of PFASs on soil structure and microbially-driven processes are currently lacking. This study explored the effects of perfluorooctanesulfonic acid (PFOS), perfluorooctanoic acid (PFOA) and perfluorobutanesulfonic acid (PFBS) at environmental-relevant nominal concentrations (1 ~ 1000 ng g^−1^) on soil functions, using a 6-week microcosm experiment. We measured soil respiration, litter decomposition, enzyme and microbial activities, soil aggregates, and bacterial abundance. PFAS (even at 1 ng g^−1^ for PFBS) significantly increased litter decomposition, associated with positive effects on bacterial abundance, and β-glucosidase activities. This effect increased with PFAS concentrations. Soil respiration was significantly inhibited by PFAS in the 3^rd^ week, while this effect was more variable in week 6. Water-stable aggregates were negatively affected by PFOS and PFOA, possibly related to microbial shifts. The general microbial activities and β-D-cellobiosidase and phosphatase activities were barely affected by PFAS treatments. Our work highlights the potential effects of PFAS on soil health, and we argue that this substance class could be a factor of environmental change of potentially broad relevance in terrestrial ecosystem functioning.

**Synopsis:** PFAS are likely to affect soil health.

**Abstract Art:** 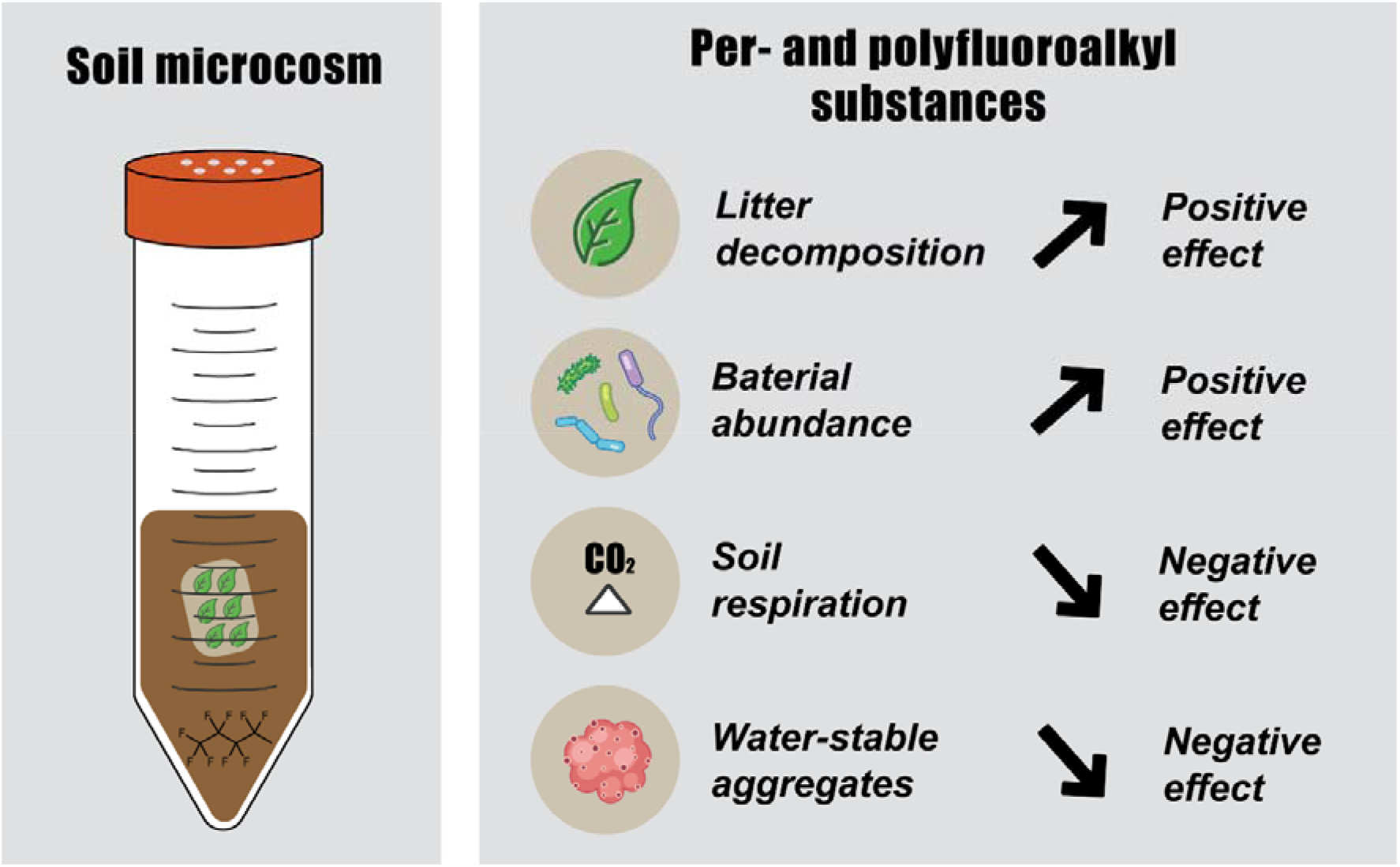

## Introduction

Human activity is progressively and fundamentally affecting the Earth’s surface, including soils, which operate at the interface between biosphere, hydrosphere, atmosphere, and lithosphere, and are suffering from physical, chemical, and biological stressors related to anthropogenic activities ^1,2^. One significant category of these influences is chemical pollution, which is widely recognized as a global change factor ^1,3^. Perfluoroalkyl and polyfluoroalkyl substances (PFAS) are a highly diverse family of chemicals of concern ^4^. These compounds contain the perfluoroalkyl moiety (C_n_F_2n+1_–) ^5^, and the relatively high-energy carbon-fluorine (C-F) bonds make them extremely resistant to breakdown, and subsequently persistent in the environment. Therefore, they are also called “forever chemicals” in public discourse ^6^. Certain PFAS, perfluorooctanesulfonic acid (PFOS) and perfluorooctanoic acid (PFOA) are on the list and on the waiting list under the Stockholm Convention ^7^, respectively.

Soil is a major sink for persistent organic chemicals in the environment. There are many pathways for PFAS entering the soil environment. Typically, fluoride factory emission, sludge application, the degradation of aqueous film-forming foam, and landfills contribute direct sources, and atmospheric deposition and runoff constitute non-point sources ^8,9^. PFAS have been widely detected in soils with a broad range of concentrations. Generally, PFAS concentrations in non-hotspot soil are lower than 300 ng g^−1^, while in hotspots of PFAS-contaminated soil, they can be as high as several or even tens of μg g^−1^, mostly dominated by PFOA and PFOS ^10^. Jin et al. ^11^ also reported that perfluorobutanesulfonic acid (PFBS) was a prominent type of PFAS in the soil around a fluoride-factory park.

Existing evidence has demonstrated that PFAS can exert some impacts on soil functions, in particular on soil enzymes, and microbial activities and communities, and these influences were closely related to properties of soil and PFASs. According to the number of carbon atoms, perfluoroalkyl carboxylic acids with 7 or more and perfluoroalkyl sulfonate with 6 or more carbons are categorized as long-chain PFAS, otherwise, short-chain PFAS ^5^. Short-chain PFBS might activate sucrase and urease activities, while long-chain PFOS might reduce these activities in soil ^12^. Cai et al. ^13^ showed that PFAS with sulfonic groups and longer chains had higher toxicity to soil microbial activities, and that soil with higher organic matter content and higher pH (neutral) exhibited lower impact by PFAS. Differences in sorption affinity of PFAS to soil are likely to regulate the impact of PFAS on soil microbes^8^. Changes in structure and function of microbial communities by PFAS were shown by previous research ^12,14–16^, and these shifts were suggested to affect soil processes and ecosystem functions. However, the implications on process rates in soil, for example, litter decomposition and soil aggregation, were not addressed.

Given the persistent nature of PFAS it is important to explore if these chemicals can affect soil process rates and properties. Here, we investigate PFAS effects on soil processes, including soil respiration, litter decomposition, soil aggregation, enzyme and microbial activities, as well as bacterial abundance. We discuss the environmental implications of our results and suggest that PFAS be considered as a global change factor of importance in terrestrial ecosystems.

## Materials and Methods

### Test soil and PFAS

The test soil was Albic Luvisol collected at the agricultural field station of Freie Universität Berlin in December 2020. The soil has a sandy loam texture (73.6% sand, 18.8% silty and 7.6% clay), with 1.87% total C, 0.12% total N and a soil pH of 7.1 ^17,18^. Fresh soil samples were thoroughly mixed, passed through a 2-mm sieve, and then stored at 4°C.

Three PFAS, namely perfluorooctanesulfonic acid (PFOS), perfluorooctanoic acid (PFOA) and perfluorobutanesulfonic acid (PFBS) were selected in this study due to their wide occurrence in the soil environment ^19^. Values of their Log *K*_ow_ are 5.26, 4.59 and 2.73, respectively ^20^, and their chemical structures and other physicochemical properties are listed in Supporting Information (SI) Table S1.

### Experimental setup

PFAS standards were dissolved in sterilized deionized water to prepare the stock solution with the concentration of 100 mg L^−1^. A five-gram portion of previously sterilized soil samples (121°C, 20 min, twice) was supplemented with appropriate doses of PFAS in solution (we used this sterilized ‘loading soil’ to avoid any exaggerated effects on soil communities ^21^), and thoroughly mixed this soil with 25 g of soil by manual stirring for 2 mins. A total of 30 g soil was placed in 50-mL mini-bioreactor tubes (Corning Inc., Corning, USA) with vented lids to establish experimental microcosms. Each tube was watered to 70% soil water holding capacity (WHC) with deionized water. The nominal concentrations of PFOS and PFOA in soil were 1, 10, 100 and 1000 ng g^−1^, and that of PFBS were 0.5, 5, 50 and 500 ng g^−1^, corresponding to their environmentally-relevant levels ^10^. The analytical method and actual concentrations of PFAS in soil are reported in SI Text S1, and Table S2, respectively. For the convenience of comparison of three PFAS, we also used the nominal concentration below. Tubes were placed in a randomized fashion inside a dark temperature-controlled incubator at 20 °C for 6 weeks, and each tube was watered weekly to maintain soil moisture. This experiment ran with 10 replications of blank control (without any PFAS added, but handled exactly the same way) and 8 replications of each treatment, for a total of 106 microcosms.

### Proxies for soil health and function

We measured well-established proxies for soil health and functions, including soil respiration, litter decomposition, enzyme activities, soil aggregates, and soil bacterial abundance. Soil respiration was measured with an infrared gas analyzer (LI-6400XT, LI-COR Inc., Bad Homburg, Germany), and litter decomposition was determined by the mass loss of tea bags ^18^. Soil enzymes activities were measured, including four enzymes concerning C (β-glucosidase and β-D-1,4-cellobiosidase), N (β-1,4-N-acetyl-glucosaminidase), and P (phosphatase) cycling, and fluorescein diacetate hydrolase (FDA) representing general soil microbial activity. Water-stable aggregates, as the basic unit of soil structure, were qualified using a wet-sieving apparatus (Eijkelkamp, Giesbeek, Netherlands) with an established method ^22,23^. Soil DNA was extracted with DNeasy PowerSoil Pro Kit (QIAGEN GmbH, Germany) following the technical protocol, and we amplified using the universal primers 515F (5′-GTGCCAGCMGCCGCGGTAA-3′) and 806R (5’-GGACTACHVGGGTWTCTAAT-3’) with quantitative polymerase chain reactions (qPCR) in a CFX 96 Real-Time System (Bio-Rad Lab., Hercules, USA). For more information on measurement procedures, qPCR conditions and quality control, see SI Text S2.

### Statistical analysis

All statistical analyses and data visualization were performed in R ^24^. The effects of PFAS treatment (four concentrations per PFAS type) on soil functions were tested with a two-step method. Firstly, we calculated the 95% confidence interval (CI) of unpaired mean differences (treatment minus control) using the R package “dabestr” ^25^. This approach focuses on the effect size and its precision, and can avoid the pitfalls of significance testing. Secondly, one-way analysis of variance (ANOVA) followed by Dunnett’s test in the R package “multcomp” was implemented to compare each treatment with the control ^26^. Model residuals were checked for heteroscedasticity and normal distribution. Spearman correlations among actual concentrations of PFAS and soil structure and function were performed with the package “corrplot” ^27^. Adjusted p values by a single-step method are reported in the SI Table S3. All data used for analyses and plotting are available online ^28^, and plots were generated with the package “ggplot2” ^29^.

## Results and Discussion

### PFAS increased litter decomposition associated with bacterial abundance

Positive effects of three PFASs on litter decomposition were observed (Figure 1A), and PFBS, particularly, at all tested concentrations significantly increased the decomposition rate (*p* < 0.05, Table S3). Regardless of tested type, increasing concentrations of PFAS significantly enhanced litter decomposition (r = 0.15, *p* = 0.0221, Figure S1 and S2B). In terms of PFAS type, PFOA and PFBS resulted in significantly positive effects on litter decomposition (*p* < 0.01, Table S3), and PFBS exerted the most remarkable positive effect (Figure S1C).

**Figure 1.**
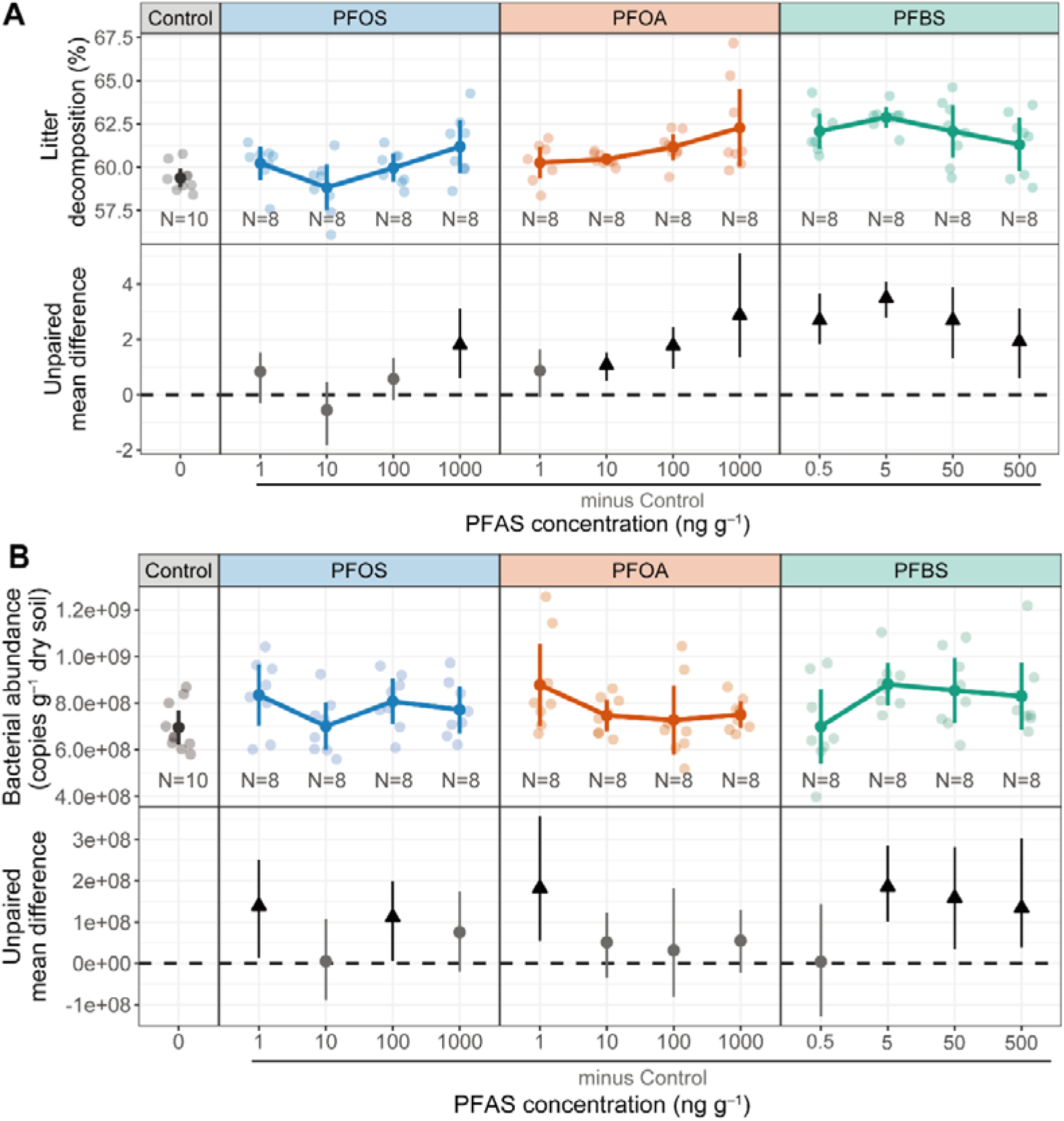
Effects of per- and polyfluoroalkyl substances (PFAS) on litter decomposition (A) and bacterial abundance (B) in soil. In the first row of each panel, raw data are presented as both scatter points and the corresponding mean and 95% confidence intervals (CIs) (N = 8 for each treatment, and N = 10 for blank control). In the second row, estimation plots present the unpaired mean difference between each treatment and the shared control. Circles in grey and triangles (arrow head up) in black represent neutral and positive effects, respectively. PFOS, perfluorooctanesulfonic acid; PFOA, perfluorooctanoic acid; PFBS, perfluorobutanesulfonic acid. Summary of effects of PFAS concentration or type refers to SI Figure S2 and S3, respectively, and outcomes of ANOVA followed by Dunnett’s test is presented in SI Table S3.

The treatment of PFASs significantly increased soil bacterial abundance (F = 1.868, *p* = 0.0485), although the multiple comparison tests did not reveal a significant difference (Table S3). PFBS seems to have the most obvious effect among three PFAS, showing a positive effect with concentrations higher than 0.5 ng g^−1^ (Figure 1B). Independently of concentration, PFAS treatments had a positive effect on soil bacterial abundance in terms of copy number per gram of dry soil (Figure S3C).

Microorganisms, including bacteria and fungi, play an essential role in the biological decomposition of organic matter in soil, a process in which bacteria generally predominate in neutral or alkaline soils, while fungi are more important in acidic soil ^30,31^. Correspondingly, we found that in our pH-neutral soil, with various PFAS treatments, the number of bacteria increased compared to the control (Figure 1B), and there was a trend for a correlation between bacteria abundance and litter decomposition rate (F=2.705, *p* = 0.103). The increased bacterial populations probably contributed to the increased decomposition rate of organic matter.

Litter decomposition and the ensuing nutrient release, governing carbon and nutrient cycling, is a key process in terrestrial ecosystems. Our results showed that PFAS had a positive effect on litter decomposition, and particularly the PFBS treatment, even at 1 ng g^−1^ in soil, resulting in a significant enhancement of litter decomposition, likely associated with increased soil bacterial abundance.

### Soil respiration is inhibited by PFAS

We observed that the tested PFAS produced significantly negative effects on soil respiration in week 3 (Figure 2A and S4), while more variable effects were present in week 6 (Figure S5). In week 3, PFOA and PFBS at all tested concentrations exerted negative effects on soil respiration, while effects of PFOS were dependent on its concentration (*F* = 5.461, *p* < 0.001). Irrespective of concentration, our tested PFAS had negative effects on soil respiration in week 3 (Figure S4C). Compared with soil respiration in week 3, less pronounced effects were observed in week 6. Similar to effects in week 3, PFBS had the most apparent effect on respiration in week 6 (Figure S5C). However, this negative effect was significant only at its highest concentration (500 ng g^−1^, *p* = 0.019, Table S3).

**Figure 2.**
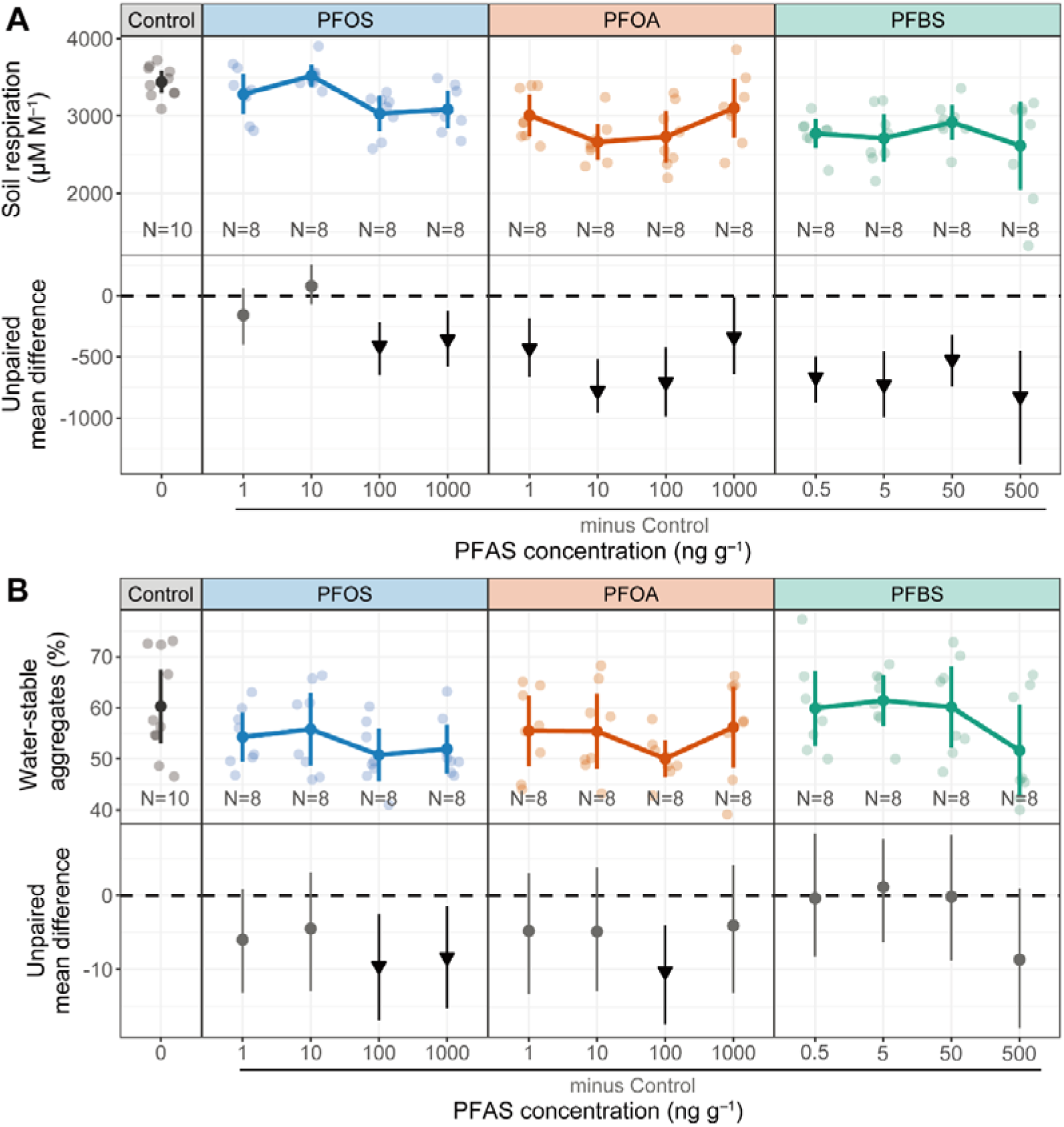
Effects of per- and polyfluoroalkyl substances (PFAS) on soil respiration at the 3^rd^ week (A) and water-stable aggregates (B). In the first row of each panel, raw data are presented as both scatter points and the corresponding mean and 95% confidence intervals (CIs) (N = 8 for each treatment, and N = 10 for blank control). In the second row, estimation plots present the unpaired mean difference between each treatment and the shared control. Circles in grey and triangles (arrow head down) in black represent neutral and negative effects, respectively. PFOS, perfluorooctanesulfonic acid; PFOA, perfluorooctanoic acid; PFBS, perfluorobutanesulfonic acid. Soil respiration at the 6^th^ week refers to SI Figure S5. Summary of effects of PFAS concentration or type refers to SI Figure S4 and S6, and outcomes of ANOVA followed by Dunnett’s test is presented in SI Table S3.

Waning effects of organic compounds on soil respiration during the incubation period have been previously reported ^32,33^. For example, the fungicides tebuconazole and carbendazim significantly suppressed soil respiration during the first 30 days, while this effect was no longer present on the 90^th^ day ^34^. Microorganisms would react to the exogenous PFAS during the first stage, expressed as an inhibitory effect, and gradually recover from this inhibition during the incubation. Another possible explanation is that as litter decomposition progressed, easily available carbon (e.g., sugars) was progressively utilized by microbes and subsequently transferred to CO_2_, which modulated the suppressed production of CO_2_.

### Limited effects on soil enzyme and microbial activities

Four enzymes were not significantly affected by individual PFAS treatments (Figures S7–S10), nor the general microbial activities (Figure S11, Table S3), but regardless of concentration, PFBS significantly increased β-glucosidase activity (*p* = 0.029, Figure S9C).

Measuring enzyme activities provides evidence on how soil biochemical processes might be affected. β-glucosidase is responsible for catalyzing the hydrolysis of cellobiose (a product of cellulose breakdown) to glucose ^35^. We observed a positive trend on β-glucosidase by PFASs, particularly PFBS, which probably contributed to the litter decomposition by PFASs. The only significant effect on β-glucosidase corresponded with the most marked effect on litter decomposition by PFBS. Additionally, there was a significant correlation between decomposition rate and β-glucosidase activity (r = 0.24, *p* = 0.004) (Figure S1).

Previous studies reported that soil dehydrogenase (proxy for total microbial activity), urease and sucrase activities were only insignificantly impacted by PFOA and PFOS with concentrations lower than 10 μg g^−1 12,36^. Cai et al.^13^ also reported that microbial activity was barely affected by PFAS at 100 μg g^−1^ in selected soils. Changes in enzyme activities are highly dynamic processes ^12,37^, and thus insignificant effects observed at harvest do not necessarily indicate that there were no remarkable changes during the incubation. In addition, there was no significant relationship between microbial activities (FDA) with other parameters (Figure S1). Overall, general microbial activities were affected only to a very limited degree.

### Different effect sizes caused by three PFAS

The three PFAS examined here appeared to exert similar impact on soil microbes and functions, but with different effect sizes, which is likely related to their bioavailability and bioaccumulation ^8^. Of the three PFASs, PFBS even at lower concentrations seemed to have the most remarkable impact on soil respiration, litter decomposition and soil bacterial abundance, while PFOS had a larger effect size on water-stable aggregates.

Sorption affinities of PFASs followed the order PFOS > PFOA > PFBS on soils with various soil textures and organic carbon contents, showing the same order of their hydrophobicity ^20^. With a low sorption affinity to soil particles, PFBS likely had an increased likelihood to interact with soil microbes, subsequently causing an impact. However, it is not a simple effect of hydrophobicity, because, for example, the higher hydrophobicity might result in higher bioaccumulation and hence exert higher toxicity in soil microorganisms ^12,13^. This might explain the more apparent effect by PFOS on some processes via soil microbes.

### Environmental implications and future perspectives

The present results highlight the potential of PFAS to induce changes in soil properties and functions. Our study comprehensively analyzed PFAS impacts on soil functions, and thus we hope that our result inspires further studies that consider the impact of PFAS on soil ecosystem functions.

Within environmentally relevant concentrations, three PFAS, especially the short-chain PFBS, had a positive effect on litter decomposition in neutral soil. This effect indicates that the PFAS present in soils now might already affect ecosystem processes. The elevated decomposition might increase the release of carbon as CO_2_, CH_4_ and dissolved organic carbon, affecting carbon sinks in soil ^38^. The precise mechanisms underpinning litter decomposition effects caused by PFAS need to be explored in further studies, with changes in microbial community composition likely playing a main role. In addition, effects are likely influenced by soil and PFAS properties.

Soil aggregation is an essential feature of soil structure, principally driven by soil biota and their interactions ^39^. Our finding that certain PFAS negatively affected water-stable aggregates could indicate far-reaching consequences for soil health, given the many influences of soil structure on virtually all soil processes. Thus, future studies might explore these effects on the soil aggregation process in greater depth, including the formation, size distribution of soil aggregates and their intrinsic connections with soil biota.

We introduce the possibility of PFAS as persistent chemicals being a potential environmental change factor. The effects of various PFAS on soil functions should now be addressed in the context of global patterns of contamination.

## Acknowledgements

BX thanks the China Scholarship Council and Deutscher Akademischer Austauschdienst (CSC-DAAD) for a postdoctoral scholarship. MCR acknowledges support from an ERC Advanced Grant (694368). We thank Daniel Lammel, Yun Liang, Tingting Zhao, and Lili Rong for their help with experimental measurements. We thank Rosolino Ingraffia for providing soil samples.

