## Supporting Information for "Effects of perfluoroalkyl and polyfluoroalkyl substances on soil structure and function"

**Text S1** Details on the determination of PFAS concentrations in soil

After harvest, the air-dried soil samples were stored at –20 °C before analysis. The chemical analysis was performed at Bundesanstalt für Materialforschung und -prüfung (BAM), Berlin, Germany. The PFAS concentration was not directly measured, but converted based on the extracted organically bound fluorine in soil. The detailed procedures are described in a previous study ^1^. The determined concentrations of PFAS are listed in Table S2.

We also found that the blank control contained tens of ng g^–1^ (Table S2). This could be the case, because we determined fluorinated organic substances in soil, but not our target PFAS, so these fluorinated organic substances can be derived from other fluorine-containing organic agrochemicals. Meanwhile, due to the relatively high content of fluorinated organic substances in the blank soil, the determination of low concentrations of PFAS (i.e., < 100 ng g^–1^) was inevitably accompanied with large errors. Even so, an apparent gradient of PFAS concentration was successfully achieved, which reached our objectives to understand soil response to gradients of PFAS concentrations.

**Text S2** Details on the measurements of the proxies of soil health and function

*Litter decomposition*

Litter bags were produced to investigate the effects of PFASs on microbial decomposition of organic matters, which is the most common approach to measure litter decomposition rate ^2,3^. A nylon mesh (30 μm) was manually cut, folded and filled with pre-weighed 300 mg commercial green tea (Sichuan Mingshan Tea Co., China). We used an impulse sealer (Mercier Corp.) to seal the litter bag up, with a size of 2.5 ×1.5 cm ^4^. Sealed bags were microwaved for 30 sec to minimize their microbial communities and buried in the middle depth of 30 g soil when placed into each tube. At the end of incubation, litter bags were picked up, rinsed by deionized water to remove adhered soil particles and dried at 60 °C. The reduced mass of green tea was employed as an indication of decomposition (%).

*Soil respiration*

Soil respiration was determined twice, after the 3- and 6-week incubation. We opened the cap of each tube for 5 mins to equilibrate the air in the tube with the ambient environment, and changed to the airtight cap with rubber stoppers. Meanwhile, 1 ml of air was sampled for the headspace of each tube as the initial point. After 4 h incubation, another 1 ml of air was sampled as the final point. Concentrations of CO_2_ in sampled air were determined by an infrared gas analyzer (LI-6400XT, LiCOR GmbH), and soil respiration was reported as the net CO_2_ production (μM M^–1^).

*Soil enzyme and microbial activities*

After incubation, a portion of 5 g fresh soil was collected to measure four functional enzymes related to C, N and P cycling, i.e., β-glucosidase and β-D-1,4-cellobiosidase (C-related), β-1,4-N-acetyl-glucosaminidase (N-related), and phosphatase (P-related). In addition, a portion of 1 g fresh soil was sampled to measure the fluorescein diacetate hydrolase (FDA) activity as an indicator of general microbial activity. Fresh soil samples were stored at 4°C before analysis, and all enzyme activities were determined in two weeks using high-throughput microplate assays (Benchmark Microplate reader, Bio-Rad Lab., Inc.). Detailed protocols were described in previous studies ^5,6^.

*Water-stable soil aggregates*

Water-stable aggregates (WSA) were quantified by the method of ^7^, as described in ^5^. In brief, 4 g dried soil was placed on a 250-μm sieve, rewetted using the capillary method, and then immersed in a wet-sieving apparatus (Eijkelkamp, Netherlands) with deionized water for 5 min. The remaining fraction on the sieve was collected and dried at 60 °C. The percentage of WSA (%) was calculated after coarse matters deducted, following WSA (%) = (remaining fraction – coarse matter) / (4.0 – coarse matter).

*Soil DNA extraction and qPCR*

We weighed approximately 250 mg fresh soil previously stored at −80 °C to extract soil DNA using DNeasy PowerSoil Pro Kit (QIAGEN GmbH, Germany), following the technical protocol. A portion DNA solution was diluted five times and stored at −20 °C for further analysis.

Quantitative polymerase chain reactions (qPCR) was operated in a CFX 96 Real-Time System (Touch 1009, Bio-Rad Lab., Inc, USA) in 96-well plates. Soil DNA was amplified with the universal primer 515F (5′-GTGCCAGCMGCCGCGGTAA-3′) and 806R (5’-GGACTACHVGGGTWTCTAAT-3’) for soil bacteria with the KAPA HiFi PCR kit (Roche GmbH, Germany). Each reaction of a volume of 20 µL contained 1×KAPA HiFi buffer, 0.3 mM KAPA dNTP Mix, 0.75 µM of each primer, 0.1 U KAPA HiFi Polymerase, 0.5 µM EvaGreen Fluorescent DNA Stain (Jena Bioscience GmbH, Germany), and 5 µL DNA template. The thermocycler was programmed with an initial denaturation step at 95 °C for 3 min, followed by 30 cycles of 95 °C denaturation for 20 s, 50 °C annealing for 30 s, and 68 °C extension for 30 s, and ended with a final extension at 68 °C for 5 min. The melting curves was tested from 65 °C to 95 °C to check the specific amplification product. All analyses were performed in triplicates, and negative controls with were also conducted in each plate. Standards for calibration were prepared from PCR amplified genes from DNA pool from soil samples using primers 27F (5’-AGAGTTTGATCMTGGCTCAG-3’), and 1492R (5’-TACGGYTACCTTGTTACGACTT-3’). The amplification product was confirmed in agarose gel for the specificity and size. After the purification with magnetic beads, the DNA concentration of PCR products was measured using a Qubit 3 Fluorometer (Fisher Scientific GmbH, Germany), and diluted to an appropriate series of points for qPCR standards. Standard curves had R^2^ values higher than 0.998, and amplification efficiencies were 89% ± 1%, in an acceptable range.

**Table S1** Basic properties of selected PFASs.

| PFAS | Purity | manufacturer | Chemical structure | Molecular formula | Molecular weight (g mol^–1^) | Water solubility | p*K*_a_ ^a^ | Log K_ow_ ^b^ |
| --- | --- | --- | --- | --- | --- | --- | --- | --- |
| Perfluorooctane sulfonic acid (PFOS) | 98% | TCI Co., Ltd. | 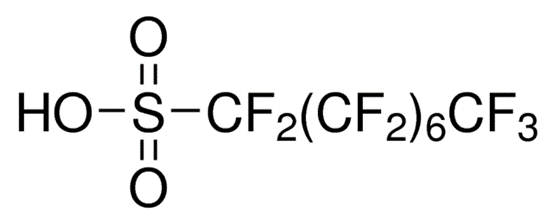 | C_8_HF_17_SO_3_ | 500.1 | Soluble ^c^ | <1.0 | 5.26 |
| Perfluorooctanoic acid (PFOA) | 98% | Strem Chemicals Inc. | 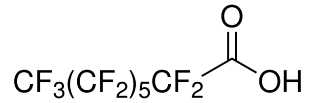 | C_8_HF_15_O_2_ | 414.0 | 3.4 g L^–1^ (20°C) ^c^ | -0.5–4.2 | 4.59 |
| Perfluorobutanesulfonic acid (PFBS) | 97% | Sigma-Aldrich Corp. | 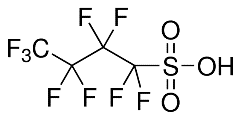 | C_4_HF_9_SO_3_ | 300.1 | 344 mg L^–1^ (25°C) ^a^ | -3.31 | 2.73 |

^a^ Data from PubChem (https://pubchem.ncbi.nlm.nih.gov)

^b^ Ref. ^8^

^c^ Data sheet from manufacturer

**Table S2** The actual concentrations of PFAS in each treatment and blank control

| PFOS (ng g^–1^) | | PFOA (ng g^–1^) | | PFBS (ng g^–1^) | |
| --- | --- | --- | --- | --- | --- |
| Nominal concentration | Actual concentration | Nominal concentration | Actual concentration | Nominal concentration | Actual concentration |
| 1 | 9.23 ± 10.09 | 1 | 4.14 ± 9.83 | 0.5 | 3.51 ± 1.71 |
| 10 | 11.48 ± 6.89 | 10 | 15.23 ± 4.37 | 5 | 6.90 ± 0.10 |
| 100 | 118.48 ± 18.19 | 100 | 110.58 ± 16.81 | 50 | 59.69 ± 0.72 |
| 1000 | 1180.57 ± 48.17 | 1000 | 1224.68 ± 142.48 | 500 | 580.10 ± 17.54 |
| Blank control | 54.45 ± 8.94 | Blank control | 118.15 ± 21.42 | Blank control | 34.86 ± 2.00 |

**Table S3** Outcomes of multiple comparison of treatments with the control for soil functions. The p-value < 0.05 was considered significant and marked in bold. In the PFAS treatment, the first column represents each concentration of four PFASs.

| **Treatment − control ≥ 0** | **Decomposition** | | **Soil respiration week 3** | | **Soil respiration week 6** | | **Water-stable aggregates** | | **Bacterial abundance** | |
| --- | --- | --- | --- | --- | --- | --- | --- | --- | --- | --- |
|  | t-value | p-value | t-value | p-value | t-value | p-value | t-value | p-value | t-value | p-value |
| **PFAS treatment** | | |  |  |  |  |  |  |  |  |
| PFOS_1 | 1.250 | 0.854 | -0.954 | 0.971 | 0.720 | 0.997 | -1.551 | 0.643 | 2.038 | 0.3065 |
| PFOS_10 | -0.828 | 0.990 | 0.484 | 1.000 | 1.057 | 0.943 | -1.163 | 0.900 | 0.073 | 1 |
| PFOS_100 | 0.850 | 0.988 | -2.470 | 0.126 | 0.887 | 0.983 | -2.462 | 0.129 | 1.636 | 0.5785 |
| PFOS_1000 | 2.679 | 0.077 | -2.154 | 0.246 | 0.263 | 1.000 | -2.160 | 0.243 | 1.112 | 0.9223 |
| PFOA_1 | 1.294 | 0.827 | -2.603 | 0.092 | -0.039 | 1.000 | -1.240 | 0.859 | 2.671 | 0.0784 |
| PFOA_10 | 1.589 | 0.615 | -4.700 | **< 0.001** | -1.587 | 0.616 | -1.263 | 0.846 | 0.752 | 0.9955 |
| PFOA_100 | 2.638 | 0.085 | -4.276 | **< 0.001** | -0.389 | 1.000 | -2.650 | 0.083 | 0.462 | 1 |
| PFOA_1000 | 4.283 | **< 0.001** | -2.046 | 0.302 | 1.494 | 0.687 | -1.057 | 0.943 | 0.806 | 0.9919 |
| PFBS_0.5 | 3.998 | **0.001** | -4.026 | **0.001** | -1.643 | 0.573 | -0.100 | 1.000 | 0.059 | 1 |
| PFBS_5 | 5.189 | **< 0.001** | -4.394 | **< 0.001** | -0.066 | 1.000 | 0.297 | 1.000 | 2.72 | 0.0694 |
| PFBS_50 | 3.991 | **0.001** | -3.158 | **0.021** | -1.633 | 0.581 | -0.041 | 1.000 | 2.324 | 0.174 |
| PFBS_500 | 2.858 | **0.049** | -4.987 | **< 0.001** | -3.193 | **0.019** | -2.242 | 0.207 | 1.964 | 0.35 |
| **PFAS type** | |  |  |  |  |  |  |  |  |  |
| PFOS | 1.197 | 0.410 | -1.572 | 0.225 | 0.918 | 0.593 | -2.366 | **0.044** | 1.532 | 0.2415 |
| PFOA | 2.971 | **0.008** | -4.205 | **<0.001** | -0.164 | 0.994 | -2.003 | 0.098 | 1.479 | 0.2644 |
| PFBS | 4.860 | **<0.001** | -5.112 | **<0.001** | -2.049 | 0.089 | -0.673 | 0.768 | 2.228 | 0.0599 |

**Table S3** Continued

| **Treatment − control ≥ 0** | **β-glucosidase** | | **β-D-cellobiosidase** | | **β-N-glucosaminidase** | | **Phosphatase** | | **FDA** | |
| --- | --- | --- | --- | --- | --- | --- | --- | --- | --- | --- |
|  | t-  value | p-  value | t-  value | p-  value | t-  value | p-  value | t-  value | p-  value | t-  value | p-  value |
| **PFAS treatment** | | |  |  |  |  |  |  |  |  |
| PFOS_1 | 0.333 | 1.000 | -1.086 | 0.932 | 0.690 | 0.998 | -0.195 | 1.000 | 0.072 | 1.000 |
| PFOS_10 | 0.946 | 0.973 | 0.351 | 1.000 | 1.792 | 0.463 | 1.101 | 0.927 | -0.700 | 0.998 |
| PFOS_100 | 1.545 | 0.648 | 0.271 | 1.000 | 2.244 | 0.206 | 0.874 | 0.985 | -0.148 | 1.000 |
| PFOS_1000 | 1.387 | 0.765 | -0.892 | 0.982 | 0.942 | 0.974 | 0.425 | 1.000 | -0.080 | 1.000 |
| PFOA_1 | 1.476 | 0.701 | 0.153 | 1.000 | 0.988 | 0.963 | 0.817 | 0.991 | 1.003 | 0.959 |
| PFOA_10 | 1.907 | 0.386 | 0.177 | 1.000 | 2.636 | 0.086 | 0.398 | 1.000 | 2.197 | 0.226 |
| PFOA_100 | 0.256 | 1.000 | 0.714 | 0.997 | 1.042 | 0.948 | -0.475 | 1.000 | 0.388 | 1.000 |
| PFOA_1000 | 0.818 | 0.991 | -0.675 | 0.998 | 1.187 | 0.888 | -0.273 | 1.000 | 1.563 | 0.634 |
| PFBS_0.5 | 2.147 | 0.250 | -0.633 | 0.999 | 0.637 | 0.999 | 0.295 | 1.000 | 0.363 | 1.000 |
| PFBS_5 | 1.858 | 0.417 | 0.347 | 1.000 | 0.166 | 1.000 | 0.220 | 1.000 | 1.169 | 0.897 |
| PFBS_50 | 1.733 | 0.506 | 0.588 | 1.000 | 1.744 | 0.498 | 1.551 | 0.644 | 0.789 | 0.993 |
| PFBS_500 | 1.827 | 0.439 | -0.413 | 1.000 | 1.309 | 0.817 | 1.197 | 0.883 | 1.483 | 0.695 |
| **PFAS type** | |  |  |  |  |  |  |  |  |  |
| PFOS | 1.409 | 0.297 | -0.449 | 0.908 | 1.857 | 0.132 | 0.732 | 0.726 | -0.286 | 0.972 |
| PFOA | 1.491 | 0.259 | 0.122 | 0.998 | 1.917 | 0.117 | 0.155 | 0.995 | 1.719 | 0.173 |
| PFBS | 2.532 | **0.029** | -0.036 | 1.000 | 1.263 | 0.372 | 1.084 | 0.480 | 1.269 | 0.369 |

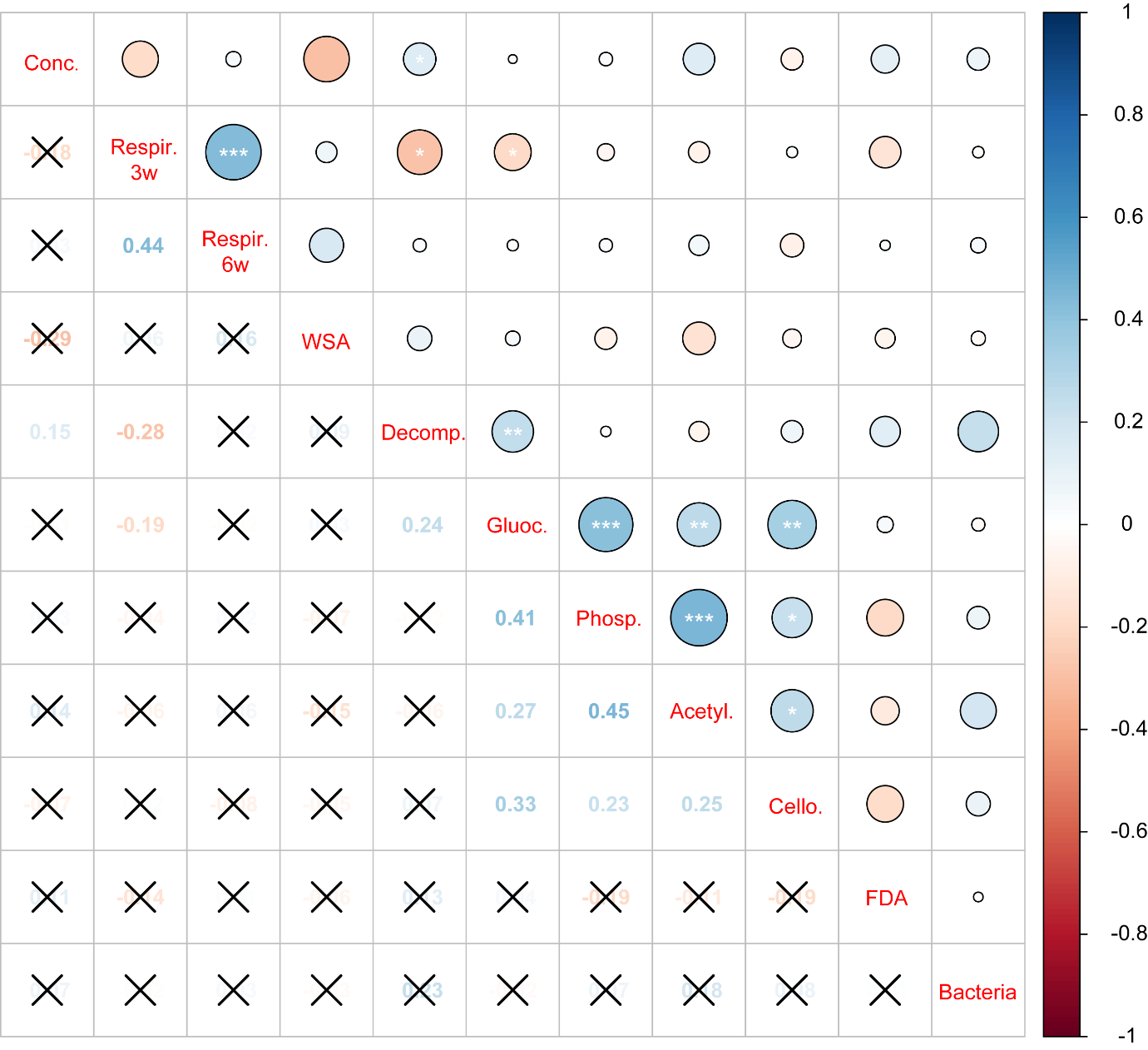

**Figure S1. Spearman correlation among actual concentrations of PFAS and proxies for soil health and function.** The significant levels of 0.05, 0.01, and 0.001 are presented as *, **, ***, respectively. Conc., PFAS concentration; Respir.3w, soil respiration at the 3^rd^ week; Respir.6w, soil respiration at the 6^th^ week; WSA, water-stable aggregates; Decomp., litter decomposition; Gluoc, β-glucosidase activity; Phosp., phosphate activity; Acetyl., β-1,4-N-acetyl-glucosaminidase activity; Cello, β-D-1,4-cellobiosidase activity; FDA, fluorescein diacetate hydrolase activity; Bacteria, soil bacterial abundance

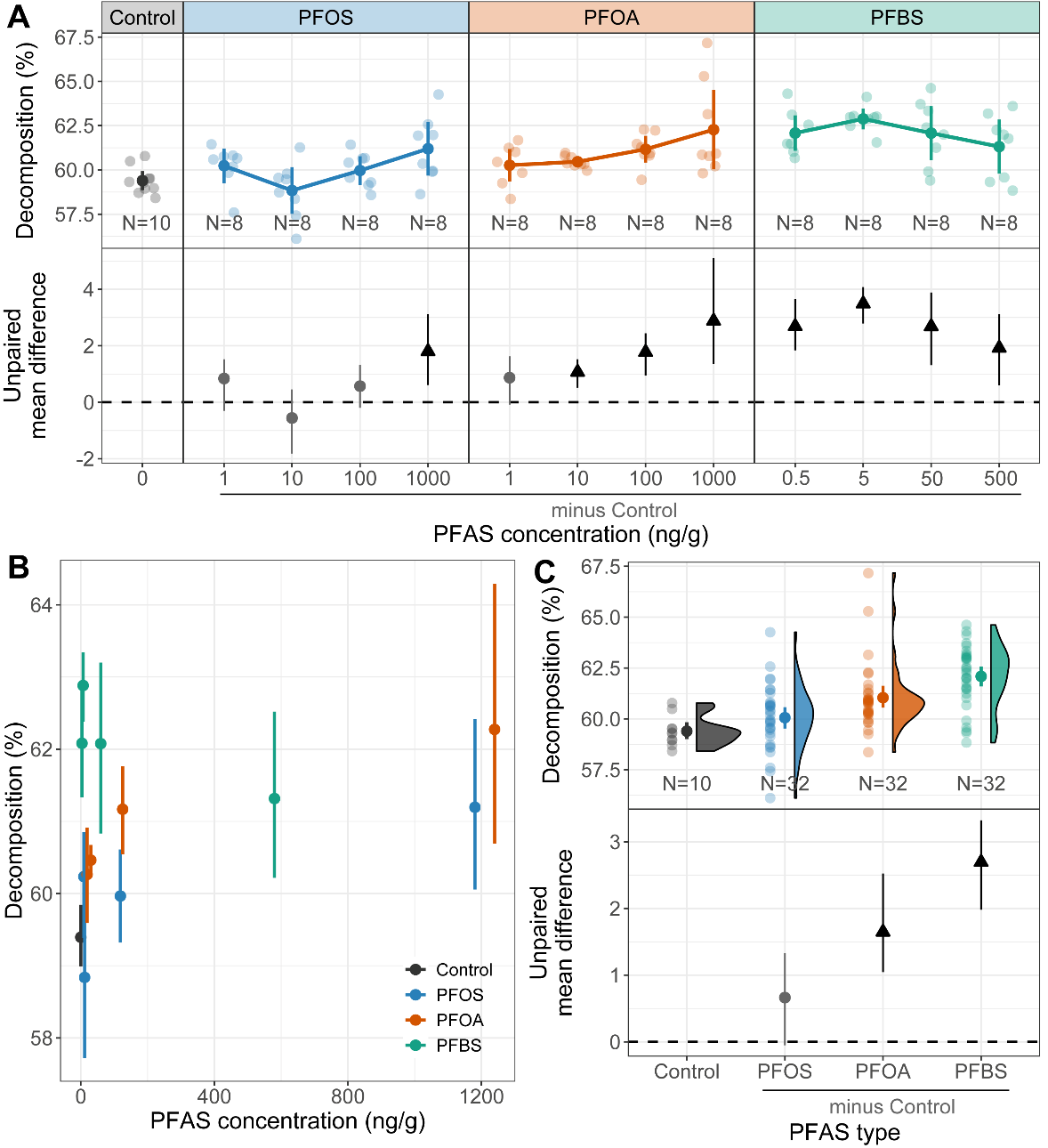

**Figure S2. Effects of per- and polyfluoroalkyl substances (PFASs) on litter decomposition in soil.** (A) Visualization of the effect of each PFAS on litter decomposition over the range of treatment concentrations (first row). Raw data are presented as both scatter points and the corresponding mean and 95% confidence intervals (CIs) (N = 8 for each treatment, and N = 10 for blank control). (B) Summary of effects of actual PFAS concentrations on litter decomposition combining treatment types. (C) Summary of PFAS type effects on litter decomposition combining treatment concentrations. Data in the first row of (C) are presented in raincloud plots supplemented with the corresponding mean and 95% CIs. In the second row, estimation plots present the unpaired mean difference between each treatment and the shared control. Circles in grey represent neutral effects (95% CIs overlapping the dashed zone line), and triangles (arrow head up) in black represent positive effects (no overlapping of 95% CIs with the dashed zero line). PFOS, perfluorooctanesulfonic acid; PFOA, perfluorooctanoic acid; PFBS, perfluorobutanesulfonic acid. The outcome of ANOVA followed by Dunnett’s test is presented in Table S3.

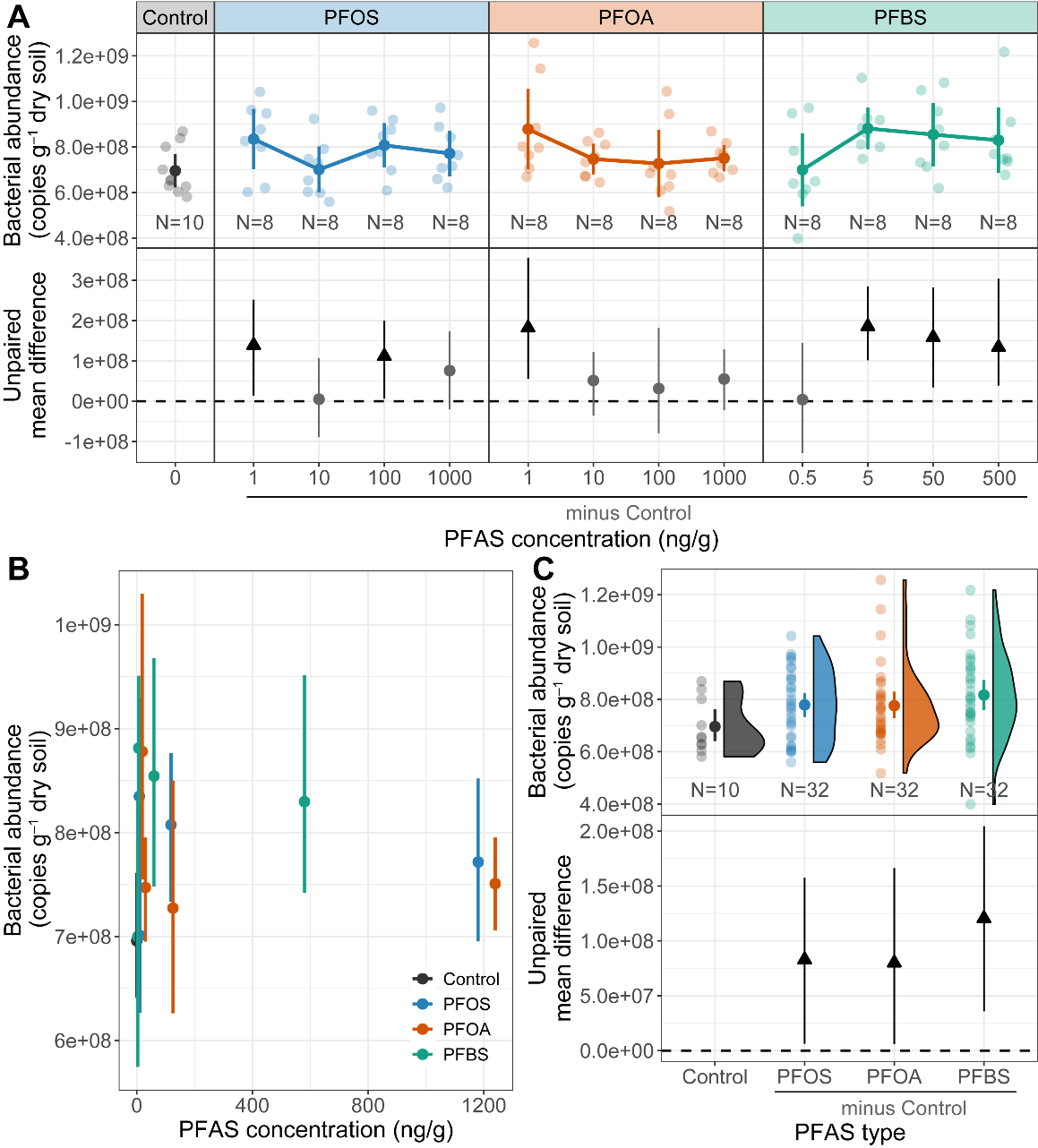

**Figure S3. Effects of per- and polyfluoroalkyl substances (PFASs) on** **soil bacterial abundance.** (A) Visualization of the effect of each PFAS on soil bacterial abundance over the range of treatment concentrations (first row). Raw data are presented as both scatter points and the corresponding mean and 95% confidence intervals (CIs) (N = 8 for each treatment, and N = 10 for blank control). (B) Summary of effects of actual PFAS concentrations on soil bacterial abundance combining treatment types. (C) Summary of PFAS type effects on soil bacterial abundance combining treatment concentrations. Data in the first row of (C) are presented in raincloud plots supplemented with the corresponding mean and 95% CIs. In the second row, estimation plots present the unpaired mean difference between each treatment and the shared control. Circles in grey represent neutral effects (95% CIs overlapping the dashed zone line), and triangles (arrow head up) in black represent positive effects (no overlapping of 95% CIs with the dashed zero line). PFOS, perfluorooctanesulfonic acid; PFOA, perfluorooctanoic acid; PFBS, perfluorobutanesulfonic acid. The outcome of ANOVA followed by Dunnett’s test is presented in Table S3.

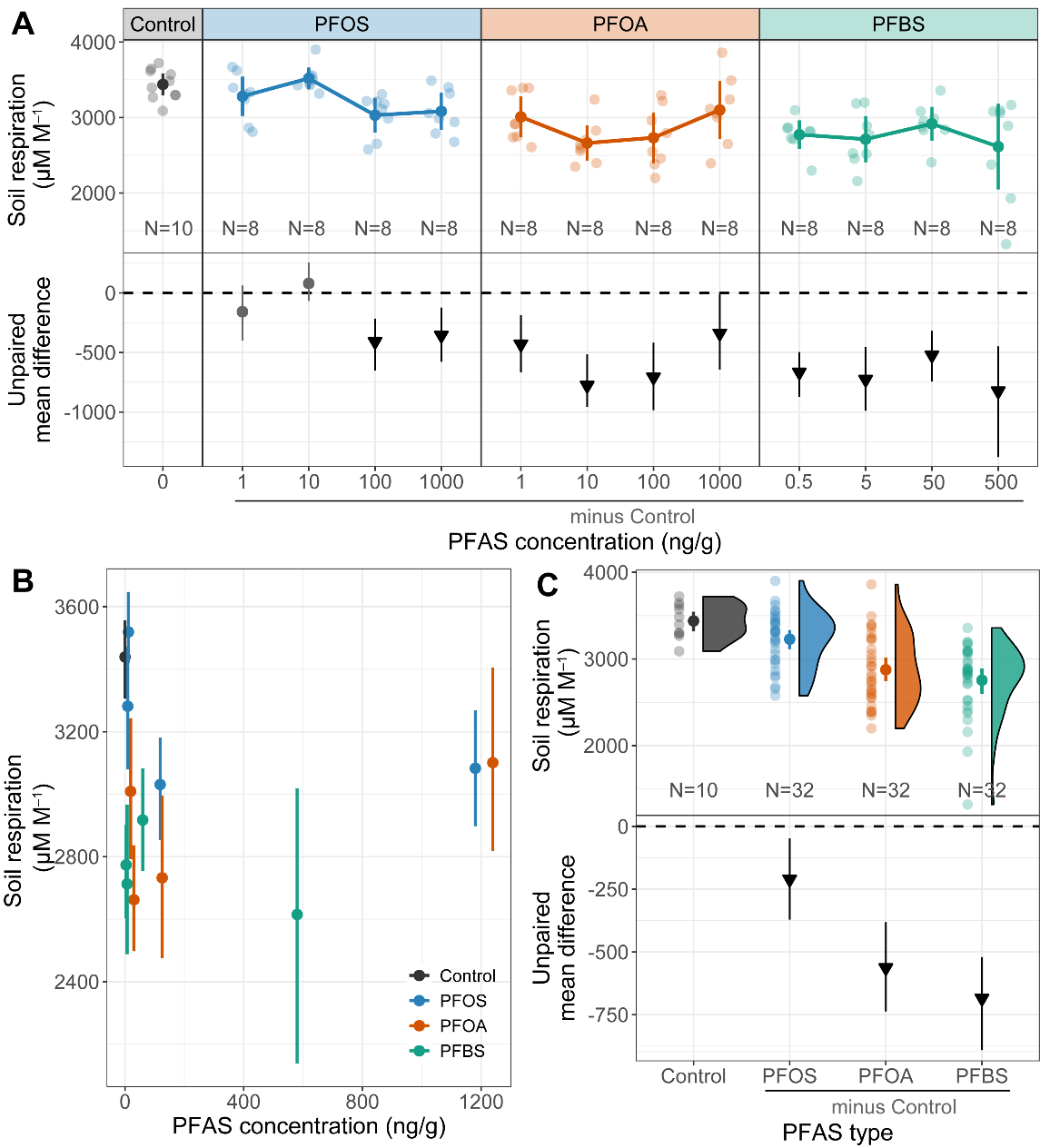

**Figure S4. Effects of per- and polyfluoroalkyl substances (PFASs) on soil respiration in week 3.** (A) Visualization of the effect of each PFAS on soil respiration over the range of treatment concentrations (first row). Raw data are presented as both scatter points and the corresponding mean and 95% confidence intervals (CIs) (N = 8 for each treatment, and N = 10 for blank control). (B) Summary of effects of actual PFAS concentrations on soil respiration combining treatment types (first row). (C) Summary of PFAS type effects on soil respiration combining treatment concentrations. Data in the first row of (C) are presented in raincloud plots supplemented with the corresponding mean and 95% CIs. In the second row, estimation plots present the unpaired mean difference between each treatment and the shared control. Circles in grey represent neutral effects (95% CIs overlapping the dashed zone line), and triangles (arrow head down) in black represent negative effects (no overlapping of 95% CIs with the dashed zero line). PFOS, perfluorooctanesulfonic acid; PFOA, perfluorooctanoic acid; PFBS, perfluorobutanesulfonic acid. The outcome of ANOVA followed by Dunnett’s test is presented in Table S3.

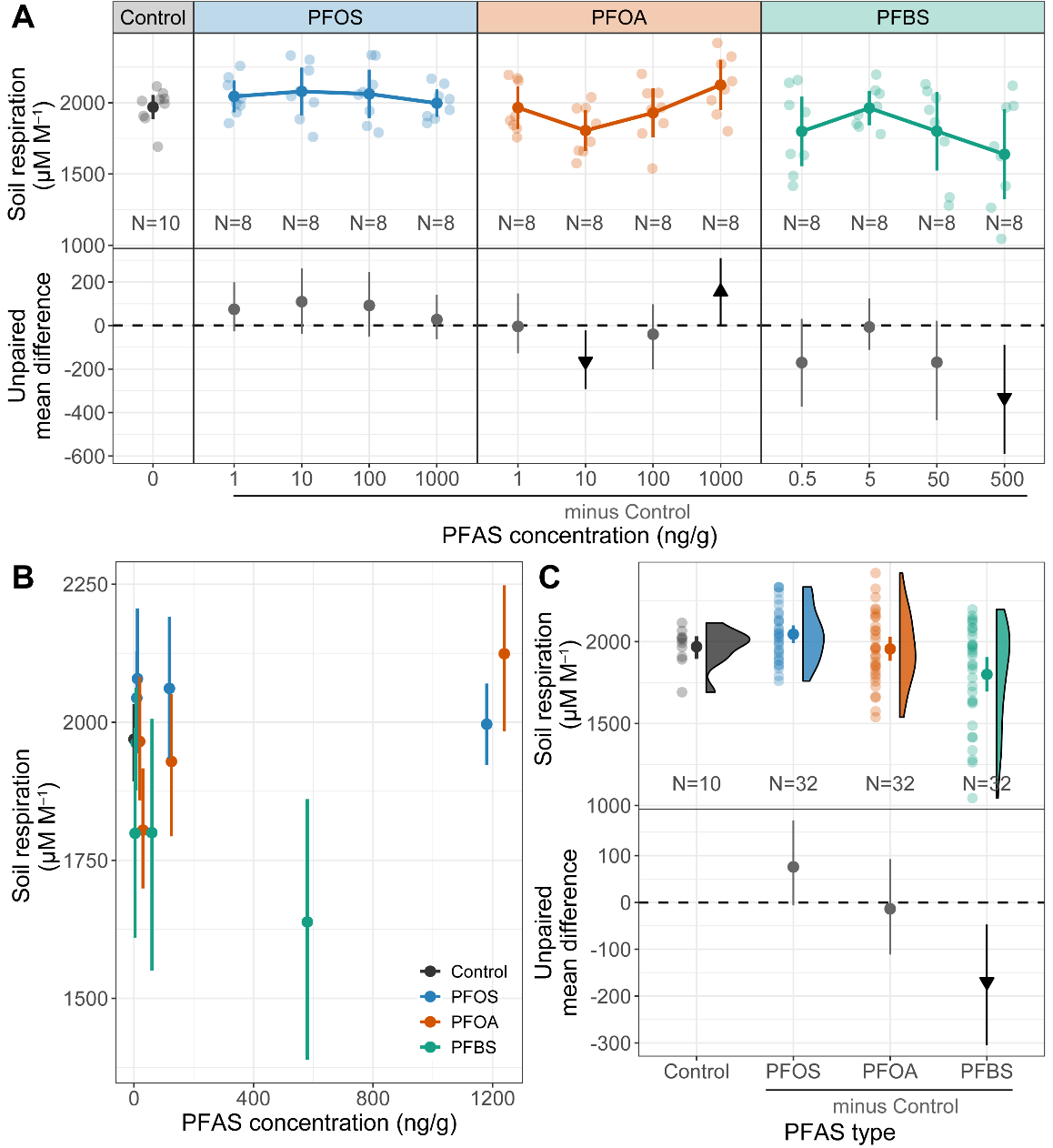

**Figure S5. Effects of per- and polyfluoroalkyl substances (PFASs) on soil respiration in week 6.** (A) Visualization of the effect of each PFAS on soil respiration over the range of treatment concentrations (first row). Raw data are presented as both scatter points and the corresponding mean and 95% confidence intervals (CIs) (N = 8 for each treatment, and N = 10 for blank control). (B) Summary of effects of actual PFAS concentrations on soil respiration combining treatment types. (C) Summary of PFAS type effects on soil respiration combining treatment concentrations (first row). Data in the first row of (C) are presented in raincloud plots supplemented with the corresponding mean and 95% CIs. In the second row, estimation plots present the unpaired mean difference between each treatment and the shared control. Circles in grey represent neutral effects (95% CIs overlapping the dashed zone line), and triangles with arrow head up and down in black represent positive and negative effects respectively (no overlapping of 95% CIs with the dashed zero line). PFOS, perfluorooctanesulfonic acid; PFOA, perfluorooctanoic acid; PFBS, perfluorobutanesulfonic acid. The outcome of ANOVA followed by Dunnett’s test is presented in Table S3.

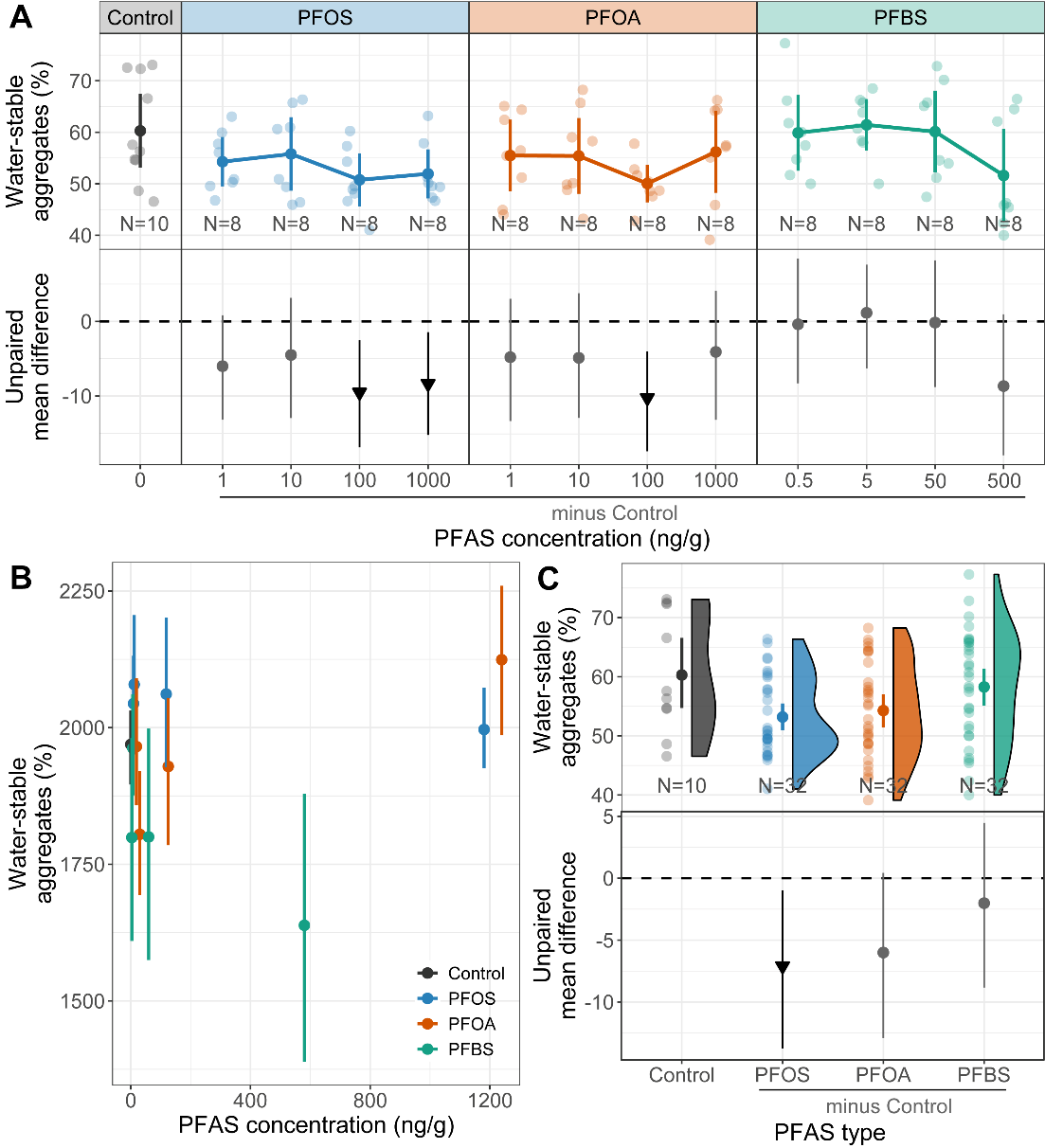

**Figure S6. Effects of per- and polyfluoroalkyl substances (PFASs) on soil** **water-stable aggregates.** (A) Visualization of the effect of each PFAS on soil water-stable aggregates over the range of treatment concentrations (first row). Raw data are presented as both scatter points and the corresponding mean and 95% confidence intervals (CIs) (N = 8 for each treatment, and N = 10 for blank control). (B) Summary of effects of actual PFAS concentrations on soil water-stable aggregates combining treatment types. (C) Summary of PFAS type effects on soil water-stable aggregates combining treatment concentrations (first row). Data in the first row of (C) are presented in raincloud plots supplemented with the corresponding mean and 95% CIs. In the second row, estimation plots present the unpaired mean difference between each treatment and the shared control. Circles in grey represent neutral effects (95% CIs overlapping the dashed zone line), and triangles with arrow head up and down in black represent positive and negative effects respectively (no overlapping of 95% CIs with the dashed zero line). PFOS, perfluorooctanesulfonic acid; PFOA, perfluorooctanoic acid; PFBS, perfluorobutanesulfonic acid. The outcome of ANOVA followed by Dunnett’s test is presented in Table S3.

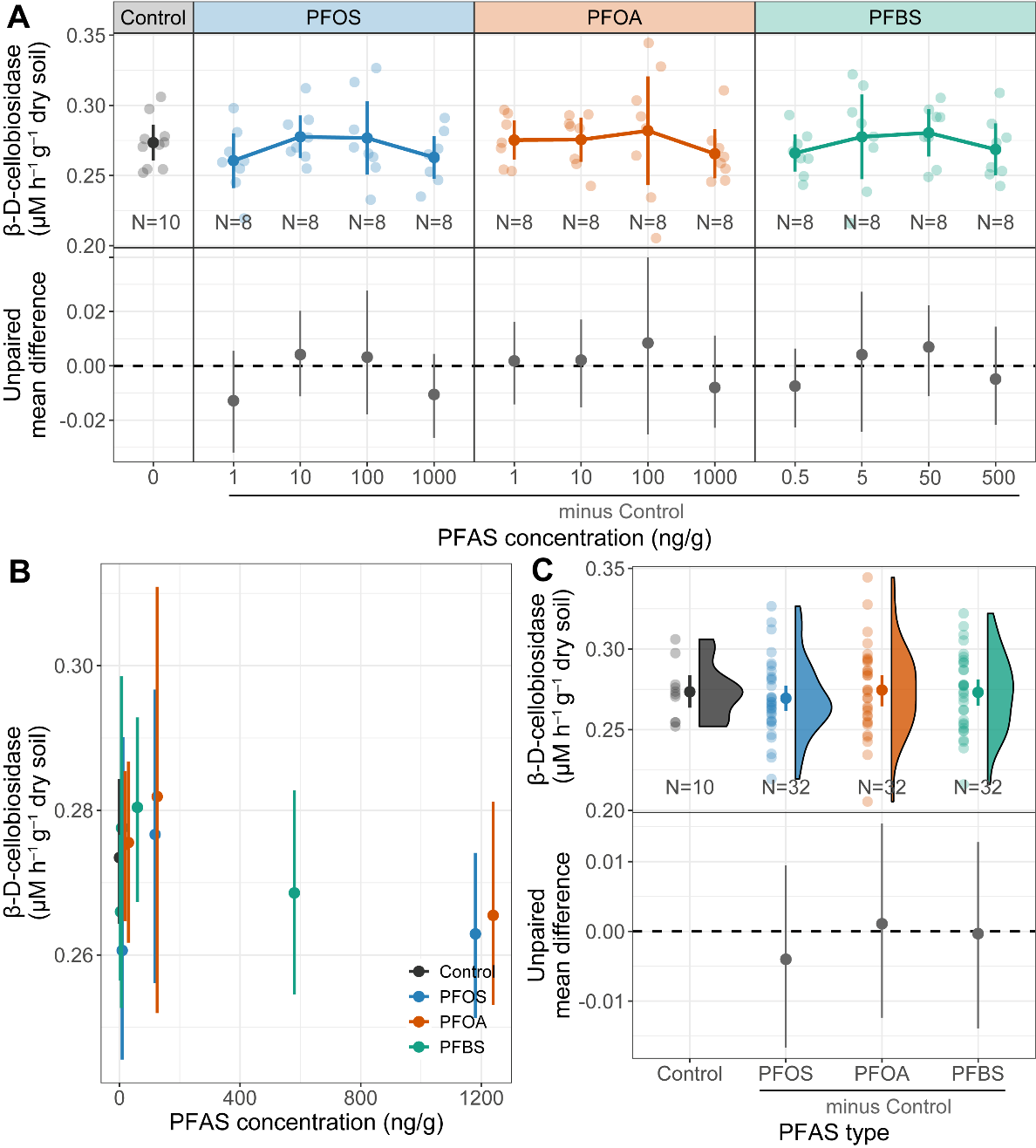

**Figure S7. Effects of per- and polyfluoroalkyl substances (PFASs) on β-D-1,4-cellobiosidase activities.** (A) Visualization of the effect of each PFAS on β-D-1,4-cellobiosidase **activities** over the range of treatment concentrations (first row). Raw data are presented as both scatter points and the corresponding mean and 95% confidence intervals (CIs) (N = 8 for each treatment, and N = 10 for blank control). (B) Summary of effects of actual PFAS concentrations on β-D-1,4-cellobiosidase activities combining treatment types. (C) Summary of PFAS type effects on β-D-1,4-cellobiosidase activities combining treatment concentrations (first row). Data in the first row of (C) are presented in raincloud plots supplemented with the corresponding mean and 95% CIs. In the second row, estimation plots present the unpaired mean difference between each treatment and the shared control. Circles in grey represent neutral effects (95% CIs overlapping the dashed zone line). PFOS, perfluorooctanesulfonic acid; PFOA, perfluorooctanoic acid; PFBS, perfluorobutanesulfonic acid. The outcome of ANOVA followed by Dunnett’s test is presented in Table S3.

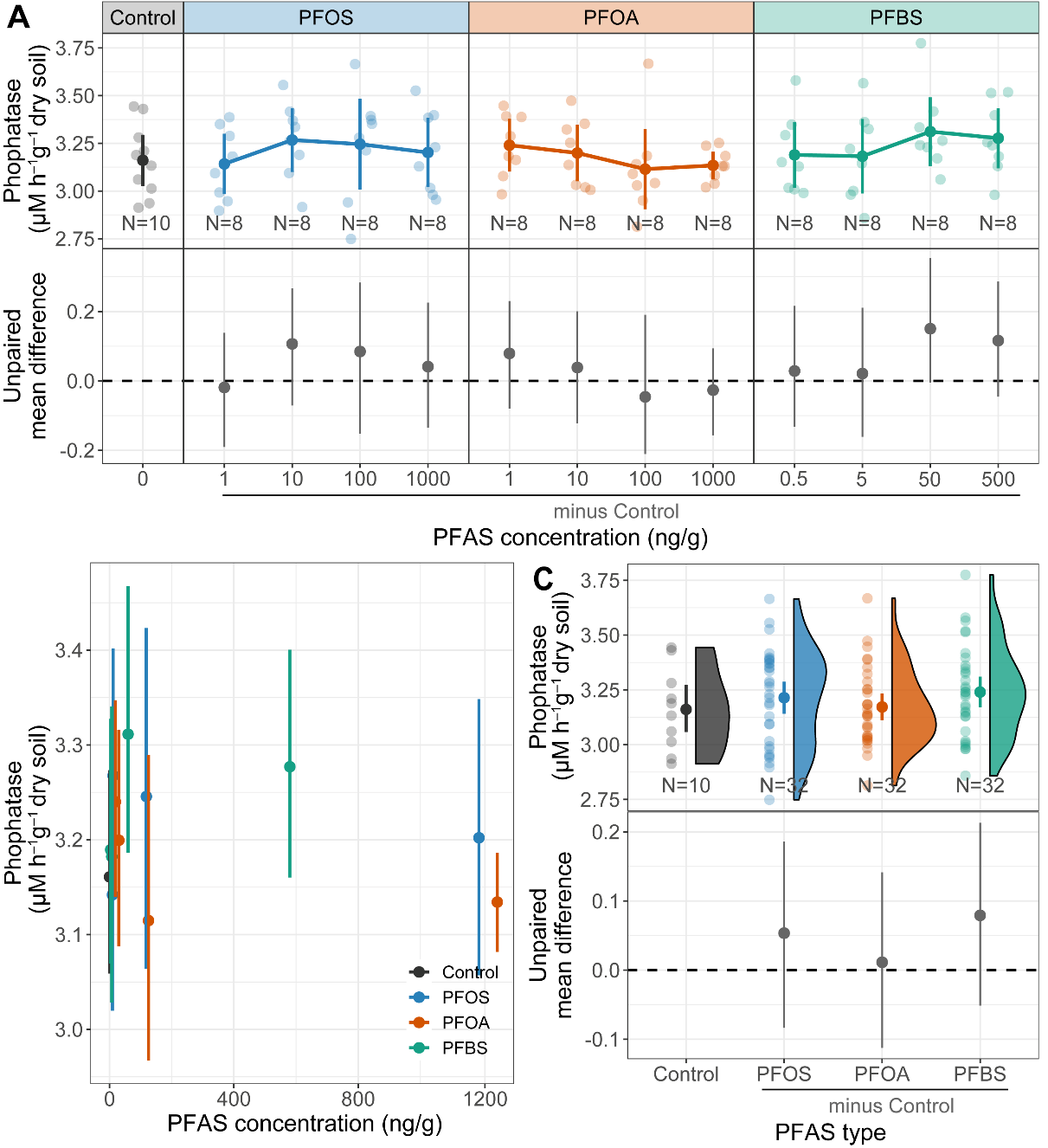

**Figure S8. Effects of per- and polyfluoroalkyl substances (PFASs) on phosphatase activities.** (A) Visualization of the effect of each PFAS on phosphatase activities over the range of treatment concentrations (first row). Raw data are presented as both scatter points and the corresponding mean and 95% confidence intervals (CIs) (N = 8 for each treatment, and N = 10 for blank control). (B) Summary of effects of actual PFAS concentrations on phosphatase activities combining treatment types. (C) Summary of PFAS type effects on phosphatase activities combining treatment concentrations (first row). Data in the first row of (C) are presented in raincloud plots supplemented with the corresponding mean and 95% CIs. In the second row of each panel, estimation plots present the unpaired mean difference between each treatment and the shared control. Circles in grey represent neutral effects (95% CIs overlapping the dashed zone line). PFOS, perfluorooctanesulfonic acid; PFOA, perfluorooctanoic acid; PFBS, perfluorobutanesulfonic acid. The outcome of ANOVA followed by Dunnett’s test is presented in Table S3.

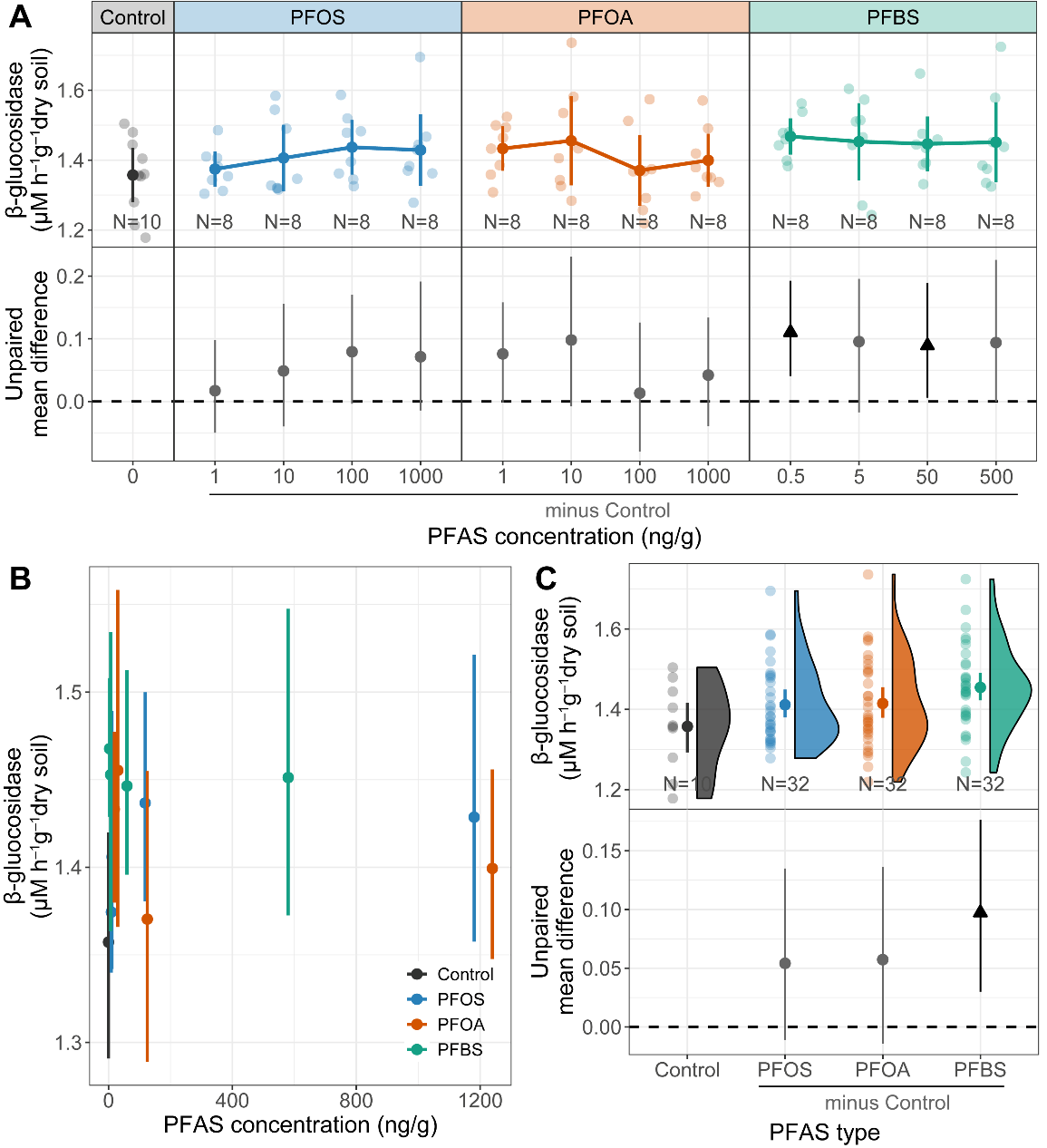

**Figure S9. Effects of per- and polyfluoroalkyl substances (PFASs) on** **β-glucosidase activities.** (A) Visualization of the effect of each PFAS on β-glucosidase activities over the range of treatment concentrations (first row). Raw data are presented as both scatter points and the corresponding mean and 95% confidence intervals (CIs) (N = 8 for each treatment, and N = 10 for blank control). (B) Summary of effects of actual PFAS concentrations on β-glucosidase activities combining treatment types. (C) Summary of PFAS type effects on β-glucosidase activities combining treatment concentrations (first row). Data in the first row of (C) are presented in raincloud plots supplemented with the corresponding mean and 95% CIs. In the second row, estimation plots present the unpaired mean difference between each treatment and the shared control. Circles in grey represent neutral effects (95% CIs overlapping the dashed zone line), and triangles (arrow head up) in black represent positive effects (no overlapping of 95% CIs with the dashed zero line). PFOS, perfluorooctanesulfonic acid; PFOA, perfluorooctanoic acid; PFBS, perfluorobutanesulfonic acid. The outcome of ANOVA followed by Dunnett’s test is presented in Table S3.

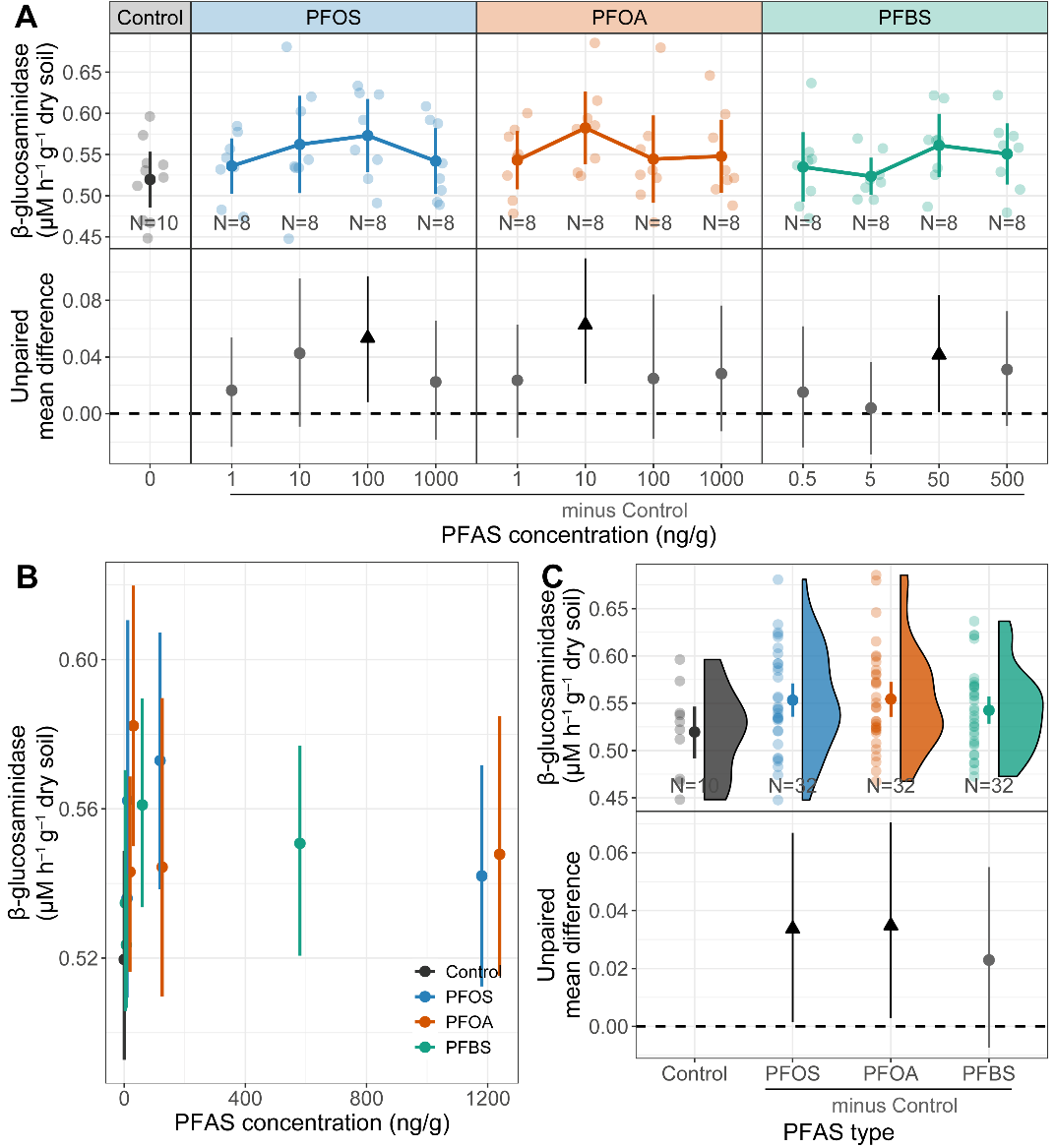

**Figure S10. Effects of per- and polyfluoroalkyl substances (PFASs) on β-1,4-N-acetyl-glucosaminidase activities.** (A) Visualization of the effect of each PFAS on β-1,4-N-acetyl-glucosaminidase activities over the range of treatment concentrations (first row). Raw data are presented as both scatter points and the corresponding mean and 95% confidence intervals (CIs) (N = 8 for each treatment, and N = 10 for blank control). (B) Summary of effects of actual PFAS concentrations on β-1,4-N-acetyl-glucosaminidase activities combining treatment types. (C) Summary of PFAS type effects on β-1,4-N-acetyl-glucosaminidase activities combining treatment concentrations (first row). Data in the first row of (C) are presented in raincloud plots supplemented with the corresponding mean and 95% CIs. In the second row of each panel, estimation plots present the unpaired mean difference between each treatment and the shared control. Circles in grey represent neutral effects (95% CIs overlapping the dashed zone line), and triangles (arrow head up) in black represent positive effects (no overlapping of 95% CIs with the dashed zero line). PFOS, perfluorooctanesulfonic acid; PFOA, perfluorooctanoic acid; PFBS, perfluorobutanesulfonic acid. The outcome of ANOVA followed by Dunnett’s test is presented in Table S3.

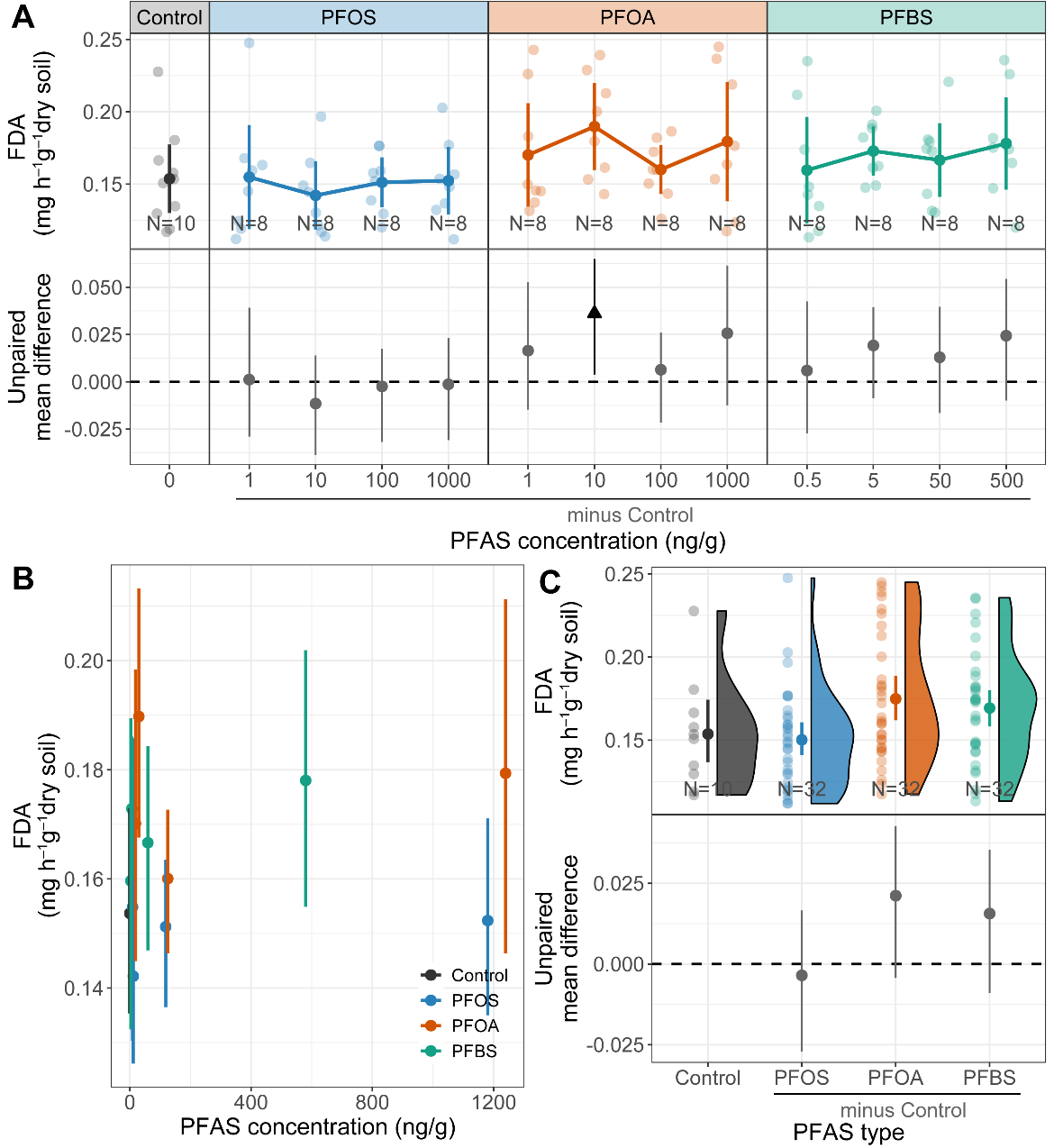

**Figure S11. Effects of per- and polyfluoroalkyl substances (PFASs) on FDA activities.** (A) Visualization of the effect of each PFAS on FDA activities over the range of treatment concentrations (first row). Raw data are presented as both scatter points and the corresponding mean and 95% confidence intervals (CIs) (N = 8 for each treatment, and N = 10 for blank control). (B) Summary of effects of actual PFAS concentrations on FDA activities combining treatment types. (C) Summary of PFAS type effects on FDA activities combining treatment concentrations (first row). Data in the first row of (C) are presented in raincloud plots supplemented with the corresponding mean and 95% CIs. In the second row, estimation plots present the unpaired mean difference between each treatment and the shared control. Circles in grey represent neutral effects (95% CIs overlapping the dashed zone line), and triangles (arrow head up) in black represent positive effects (no overlapping of 95% CIs with the dashed zero line). PFOS, perfluorooctanesulfonic acid; PFOA, perfluorooctanoic acid; PFBS, perfluorobutanesulfonic acid. The outcome of ANOVA followed by Dunnett’s test is presented in Table S3.
